# EM-LDDMM for 3D to 2D registration

**DOI:** 10.1101/604405

**Authors:** Daniel Tward, Michael Miller

## Abstract

We examine the problem of mapping dense 3D atlases onto censored, sparsely sampled 2D target sections at micron and meso scales. We introduce a new class of large deformation diffeomorphic metric mapping (LD-DMM) algorithms for generating dense atlas correspondences onto sparse 2D samples by introducing a field of hidden variables which must be estimated representing a large class of target image uncertainties including (i) unknown parameters representing cross stain contrasts, (ii) censoring of tissue due to localized measurements of target subvolumes and (iii) sparse sampling of target tissue sections. For prediction of the hidden fields we introduce the generalized expectation-maximization algorithm (EM) for which the E-step calculates the conditional mean of the hidden variates simultaneously combined with the diffeomorphic correspondences between atlas and target coordinate systems. The algorithm is run to fixed points guaranteeing estimators satisfy the necessary maximizer conditions when interpreted as likelihood estimators. The dense mapping is an injective correspondence to the sparse targets implying all of the 3D variations are performed only on the atlas side with variation in the targets only 2D manipulations.

## 1 Introduction

The field of connectomics is combining high resolution neuroimaging with big data analytics to transform our understanding of neuroscience[1]. Connectivity is reliably estimated from tracing experiments, with images acquired at micron resolution. Macroscopic techniques such as diffusion tensor imaging (DTI)[2] (which estimates structural connectivity via tractogrophy [3]), or functional magnetic resonance imaging (which estimates functional connectivity via blood oxygen level dependent (BOLD) signal correlation [4]) are merely proxies and can often be inaccurate [5].

While some micron scale 3D imaging modalities are becoming available, such as CLARITY[6, 7] and iDISCO [8], it remains that 2D histological preparations consisting of many sections which sparsely sample the true 3D volume of the brains continues to be ubiquitous to many biomedical science laboratories studying brain tissue at meso and micron scales.

For the community to benefit from this data, alignment to common atlas coordinates such as the Allen Common Coordinate Framework (CCF) [9] is essential. This technique allows for interpretation of data within an anatomical ontology such as that specified in the Allen Reference Atlas[10], and enables building statistical ensembles among different animals, as well as collaboration between different labs. One such collaborative group is the BICCN https://www.braininitiative.nih.gov/brain-programs/cell-census-network-biccn, whose goal is to “generate comprehensive 3D common reference brain cell atlases that will integrate molecular, anatomical, and functional data for describing cell types in mouse, human, and non-human primate brains”. The challenge therefore remains that complete atlases of the brains are dense 3-dimensional (3D) data sets, yet analysis technologies associated to sparse 2-dimensional (2D) sampling is still essential.

When considering how to sample this data, a dense set of slices can enable accurate 3D registration to an atlas [11]. On the other hand, imaging fewer slices saves experimental time and effort, and allows tissue to be saved for other purposes such as the multiple markers important to the BICCN. As slices become sparser, the assumption that they are similar to their neighbors is violated, and existing automatic image registration methods cannot be used.

This paper focuses on the 3D reconstruction of a sparse set of tissue slices, guided by using a well characterized atlas. Slices may be different modalities or have missing tissue or artifacts, using the approaches described in [12]. For accommodating sparse sectioning, no assumption of similarity between neighboring slices is necessary. We exploit the random orbit model of Computational Anatomy in which the measured targets are modeled as a diffeomorphic change of coordinates of a three-dimensional atlas along with a deterministic and statistical transformation applied to the atlas to generate the data.

Our method involves a sequence of transformations describing how each 2D slice is generated from a 3D atlas, resulting in a generative statistical model for observed images. Because this setting involves joint optimization over hundreds of transformations, a scalable approach for computing transformations and their gradients is required. Optimizing over multiple transformations linked by composition is the subject of deep learning, and we borrow the notion of a computation graph from this field. In this formulation data and operations are represented by edges and vertices (respectively) of a directed graph. Data flows from parent to child under the operations, and gradients flow from child to parent using each operation's derivative adjoint (also known as backpropagation or the chain rule).

This computation graph formulation of image registration has been considered previously [13]. In this work we jointly optimize over a larger series of transformations to accommodate 3D to 2D mapping, and we precisely specify each computation and its gradient, showing a comparison to approaches derived assuming continuous functions.

In this paper we describe the formulation of our registration procedure, discussing numerical issues revealed by this procedure. We illustrate its effectiveness relative to a gold standard using simulated data of increasing sparsity, and we show its performance registering data from connectomic tracing experiments as part of the BICCN.

## 2 Methods

In this section we first describe the sequence of transformations in our model formulated in terms of continuous functions. We then turn to details of discretization and interpolation. Finally we describe our experiments.

### 2.1 Synthesis Model of Generated Data

In image analysis synthesis is a fundamental part of the analysis step, implying the generative model step is fundamental to understanding the reconstruction. The synthesis model is a sequence of transformations which compose. The generative model of imaging data is shown in Fig. 1. We follow the random orbit model from Computational Anatomy. The space of dense 3D images in the orbit of the atlas are defined via diffeomorphisms *φ*: (*x*_1_, *x*_2_, *x*_3_) ∈ ℝ^3^ ↦ *φ*(*x*) = (*φ*_1_(*x*), *φ*_2_(*x*), *φ*_3_(*x*)) ∈ ℝ^3^. The diffeomorphism 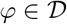 act on the atlas to generated the orbit of imagery 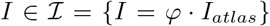, the action on the atlas defined as inverse on the right *φ* · *I* = *I* ∘ φ^-1^.

**Figure 1:**
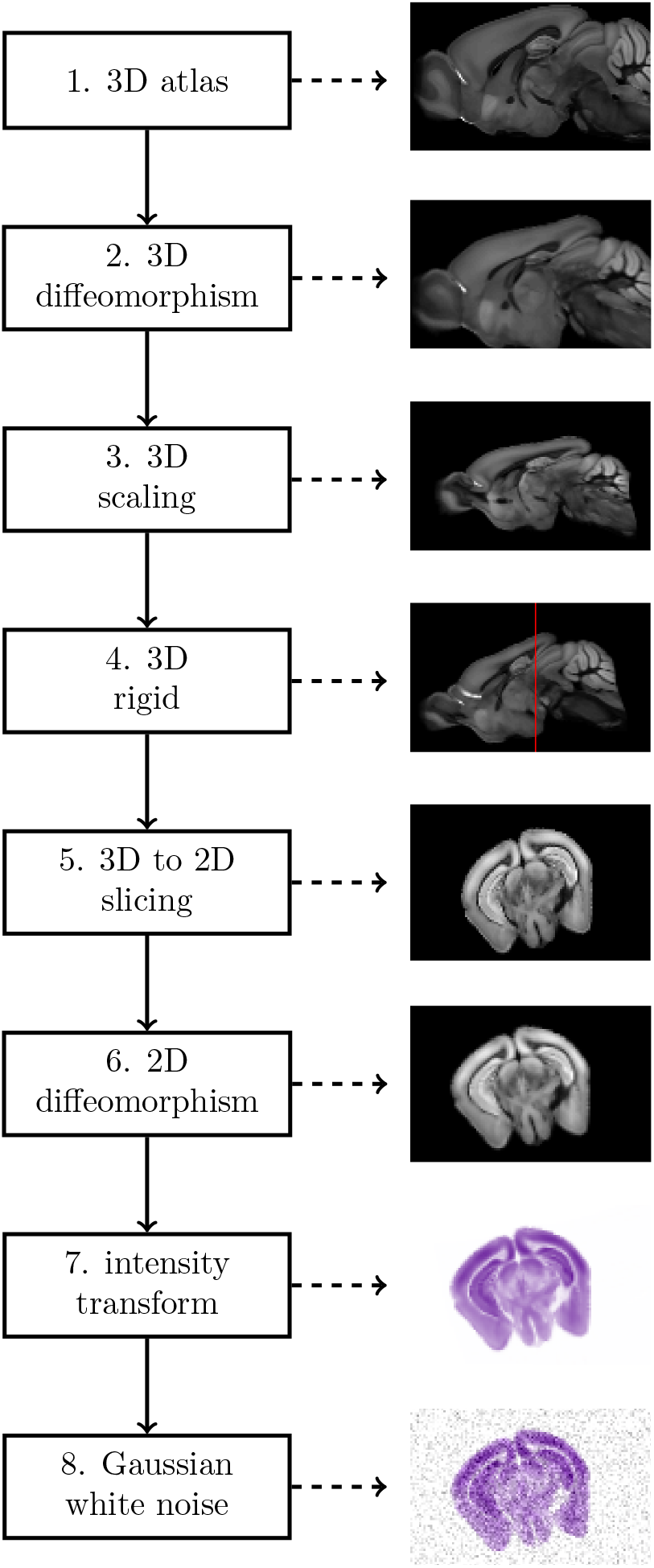
Flowchart illustrating steps in our generative model, predicting one slice of data from the Allen CCF.

We build the diffeomorphisms to correspond to a low dimensional element of the affine group *φ*_(*A,b*)_ and the infinite dimensional diffeomorphisms *φ_V_*, and they act in composition *φ* = *φ_V_* ∘ *φ_A_*. We generate them using polar decomposition and flows; see Appendix A.

The measured target data *J^i^, i* = 1,& correspond to sections that are stained and may be distorted by the sectioning and imaging process. For this we define *ψ^i^*, each *ψ^i^*: *I* → *ψ^i^*(*I*) ∈ ℝ^2^. The action of *ψ_j_* is to section through the 3D volume generating a plane, and as well to apply a diffeomorphism on the coordinates of the plane. The pointwise imaging model in the plane becomes

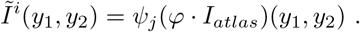

This slicing procedure is shown in Fig. 2.

**Figure 2:**
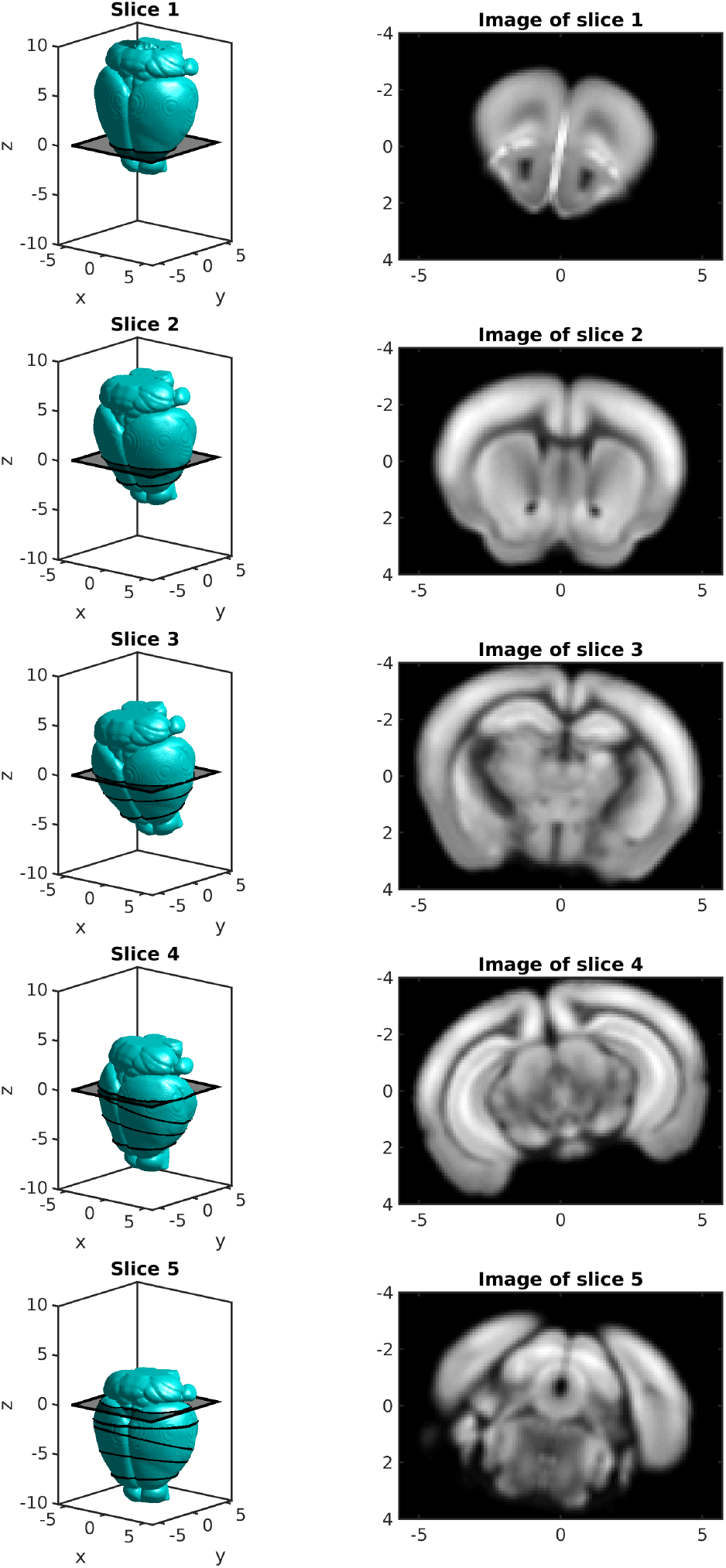
Illustration of the 3D to 2D slicing procedure. Left: 3D rigid motions are applied successively to the deformed atlas (cyan isosurface), positioning it with respect to a hypothetical cutting blade at *z* = 0. Slices need not be parallel nor uniformly spaced. Right: Sampling the rigidly positioned atlas image at *z* = 0 yields a 2D slice image.

We model our imaging system as an unknown polynomial intensity and color transformation of this sliced atlas, 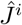, with additive white Gaussian noise.

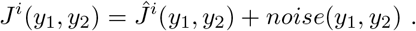

Missing tissue or artifacts are incorporated via Gaussian mixture modeling at each pixel, as described in [12]. This model corresponds to a penalized likelihood optimization algorithm that minimizes weighted sum of square error with weight *W^i^*(*y*_1_,*y*_2_).

### 2.2. Discrete formulation

The method is illustrated using a computation graph in Fig. 3, where edges correspond to data and nodes correspond to operations. Note that all transfor-mations are composed and sliced before transforming the image, avoiding loss of resolution due to repeated interpolations.

**Figure 3:**
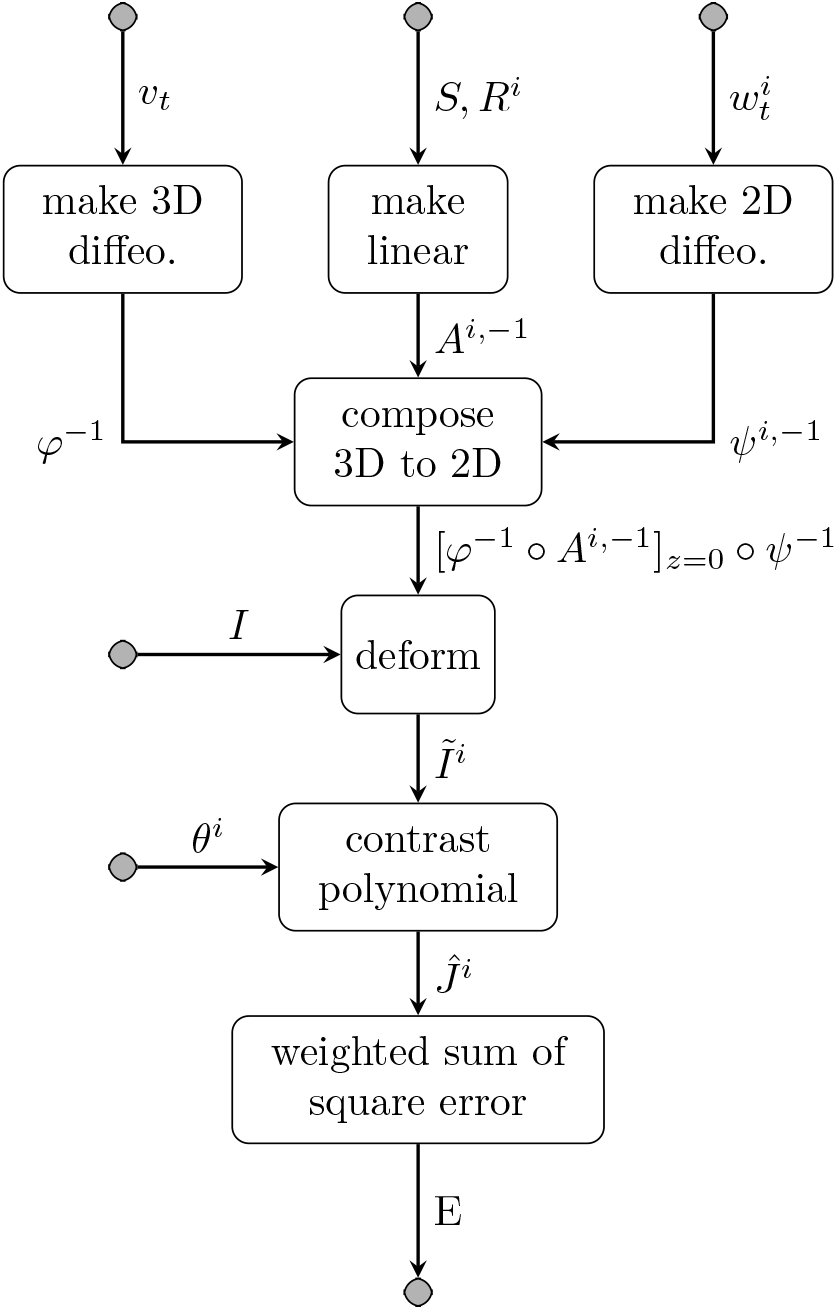
Overview of computation graph illustrating transformations for slice *i*.

A discrete version of the method is formed by specifying exactly how diffeo-morphism are generated, how transformations are composed, how the image is deformed, and how sampling grids are defined. Most of these operations are performed via linear interpolation, and are shown in Fig. 4. Operations that have more than one input argument use arrows to specify which input argument is used. For example interp ↑ (←) means “evaluate the the data specified above, at the points specified to the left, via linear interpolation.” Data defining images and vector fields are defined on grids of various sizes, which are denoted next to the edges in the graph. For example, the pixels in the *i*th observed RGB image *J^i^*, are stored on a grid with locations *X^J,i^* of size 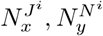, 1, 3.

**Figure 4:**
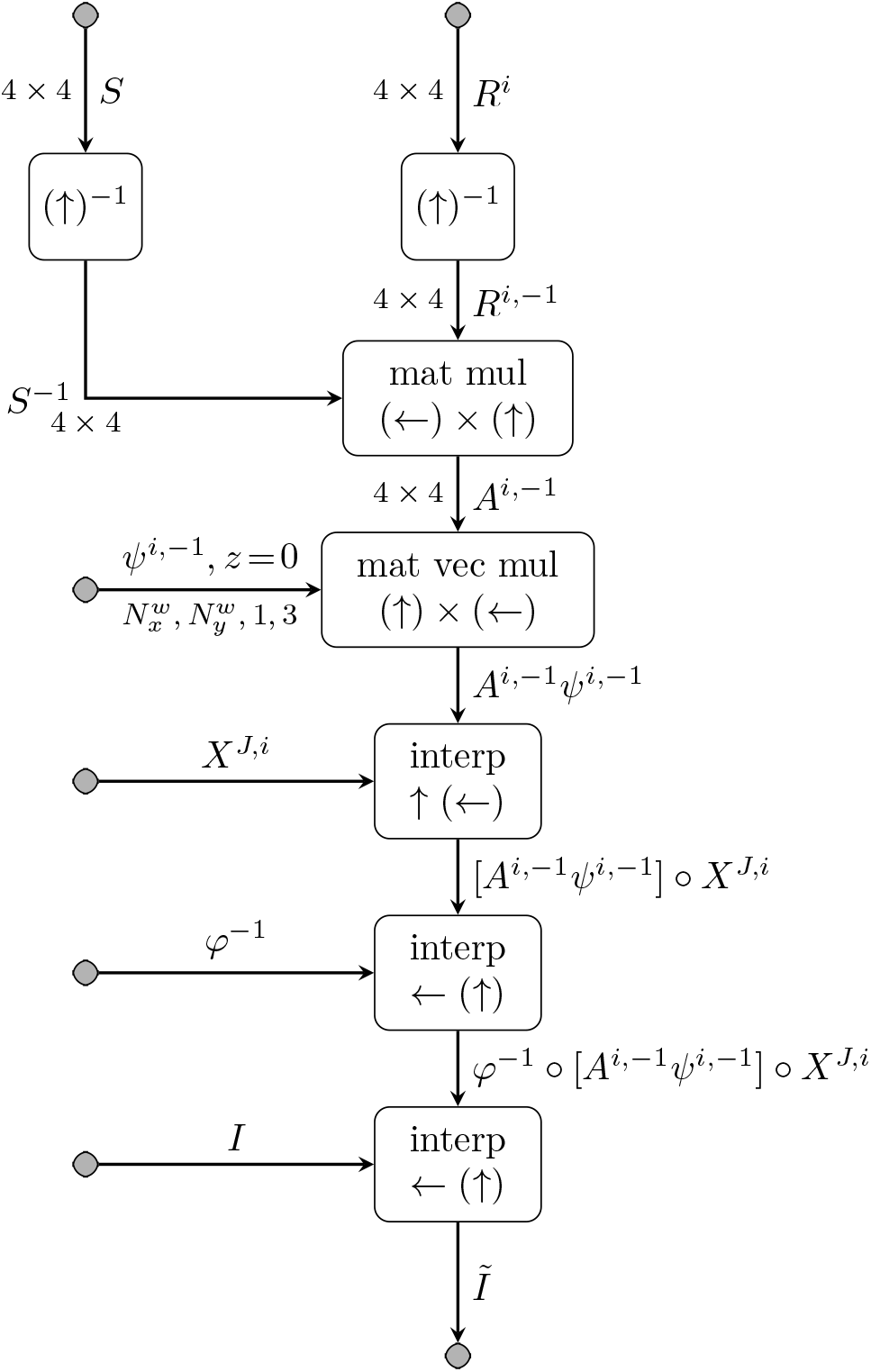
Generation of deformed image. Note that the 3D to 2D step occurs because we sample our transformations at the grid points *X^J,i^*.

Finally the integration of the velocity field *υ_t_* under optical flow to produce *φ*^-1^ is shown in Fig. 5. This is performed using linear interpolation via the method of characteristics [14].

**Figure 5:**
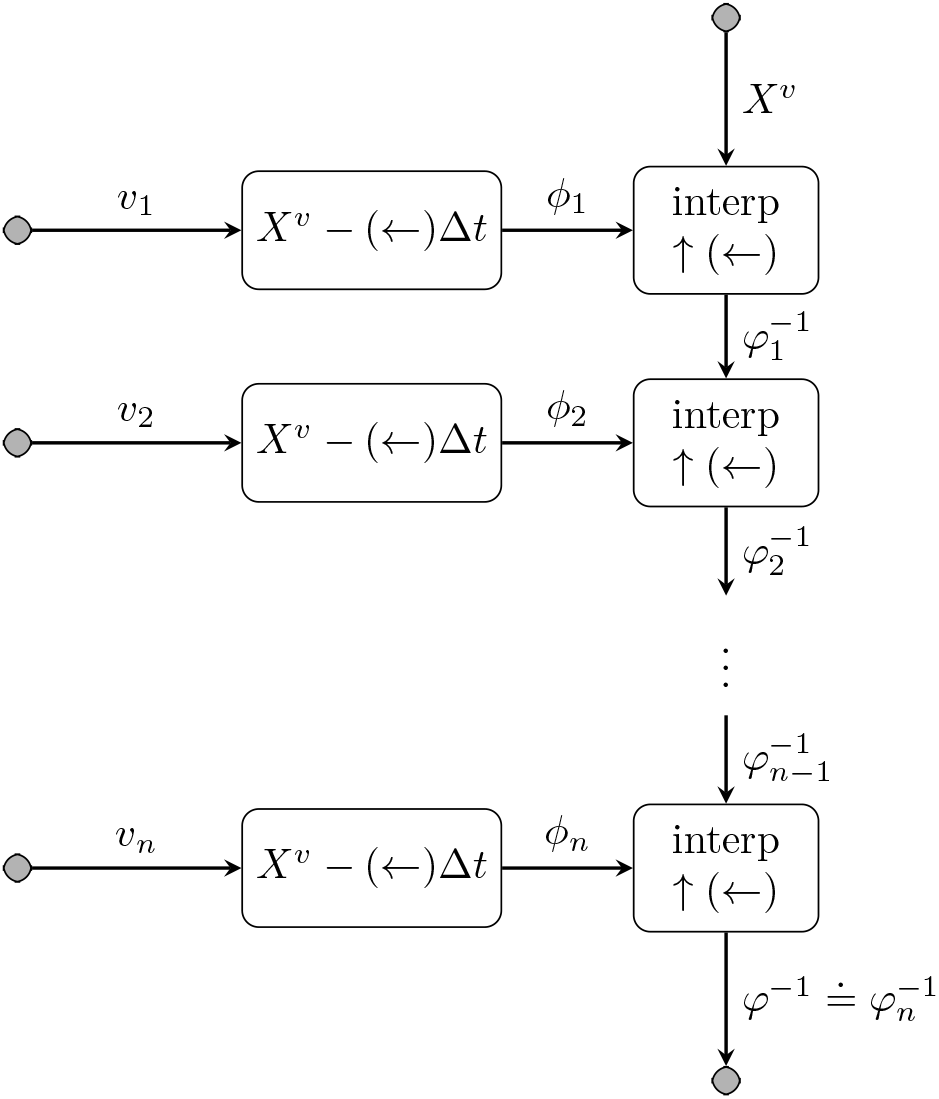
Generating 3D diffeomorphism from velocity. Small deformations are generated by subtracting displacement from identity, and they are combined under composition. This is the method of characteristics (along a characteristic curve, the value of *φ*^-1^ is constant). All data is of the same size, 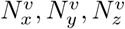, 3. Note that the 2D velocity field is generated the same way, with 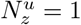.

### 2.3 Gradient calculations

For each operation, gradient backpropagation is performed using the adjoint of the operation’s derivative. That is, we define an operation *f*: *X* → *Y* with *x* ↦ *f* (*x*). Its derivative is a map between tangent spaces *Df_x_*: *T_x_X* → *T_f(x)_Y* with *δ* ↦ *Df_x_* (*δ*). The adjoint is a map between cotangent spaces, 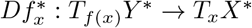 with *e* ↦ *D_x_f* *(*e*), defined implicitly by preserving the pairing

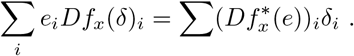

The same definition is true in the continuous case, with the sum replaced by a integral. The only two operations in our computation graph with non trivial adjoints are matrix inverse and interpolation.

#### 2.3.1 Matrix inverse

We consider the operation *f* (*A*) = *A*^-1^.

##### Theorem 2.1.

*The adjoint of the derivative of a matrix inverse, f* (*A*) = *A*^-1^, *is given by*

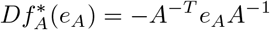

*Proof.* The action of the derivative on a perturbation *δ_A_* is given by

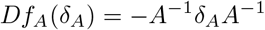

where adjacent terms denote matrix multiplication. The adjoint is calculated as

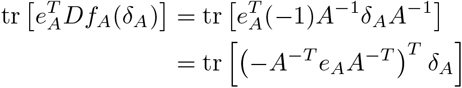

using the cyclic property of the trace.

#### 2.3.2 Discrete interpolation

In the discrete case we consider working with *ND* arrays. We transform an image 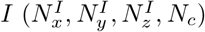 by sampling it at the points in 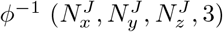. Note that 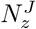 may be 1. The output *J* has size 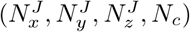.

The trilinear interpolation operation *J* = *f* (*I,φ*^−1^) is defined as

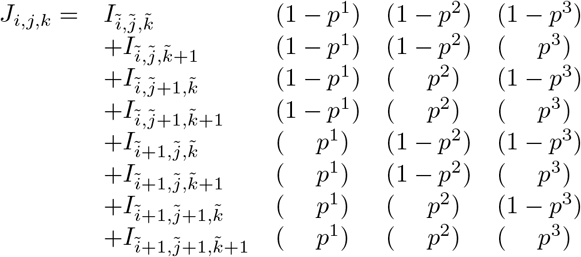

where 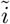 is the *i* index pointed to by 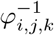, rounded down, and *p*^1^ is the fraction of the way to the next pixel.

Conceptually this can can be written as matrix multiplication for a very large sparse matrix, which can be calculated by accumulation as shown in Algorithm 1. In practice we use a more efficient implementation.

##### Algorithm 1 Calculate *J* from *I* and *φ*^−1^

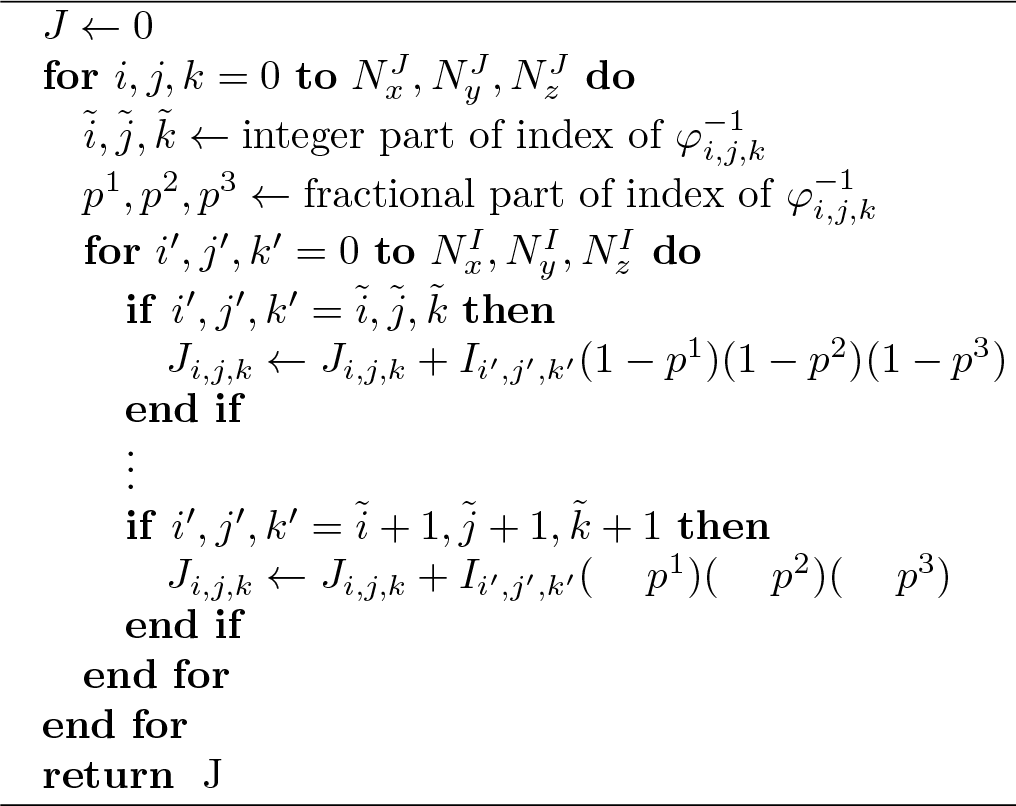

Since *f* is linear in its first argument, the adjoint is just the transpose. The adjoint can be written as shown in Algorithm 2. In practice we use a more efficient function such as Matlab’s `accumarray`.

##### Theorem 2.2.

*The adjoint of the derivative of discrete interpolation, with re-spect to its second argument is given by (for the x component)*

#### Algorithm 2 Calculate *e*^*I*^ from *e*^*J*^ and *φ*^−1^ using trilinear interpolation.

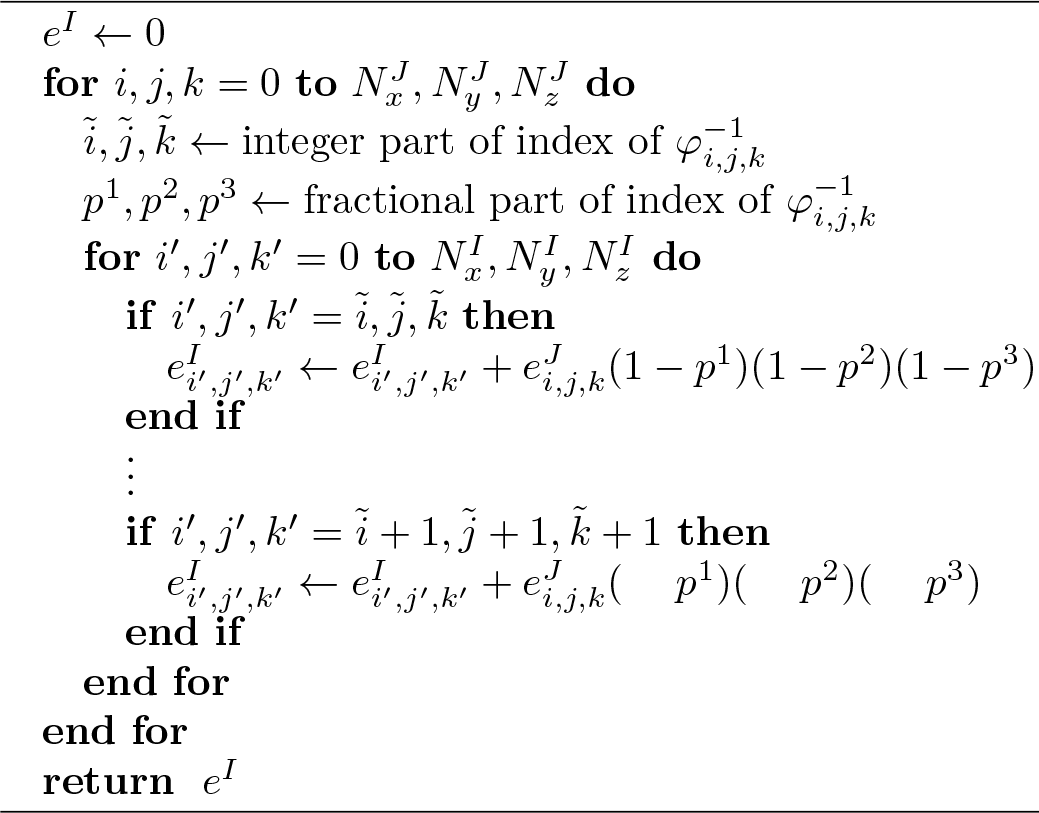

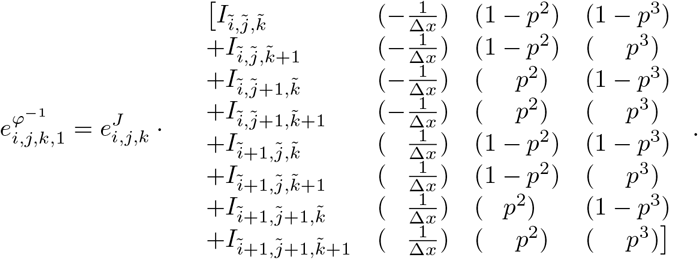

*Here 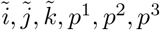 are defined as above. The adjoint is defined similarly for the y and z components.*

*Proof*. The adjoint with respect to the second argument can be computed by noticing that 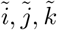 do not vary for small changes in the input argument, and *p*^1^,*p*^2^,*p*^3^ vary linearly, with slope of one over the corresponding voxel dimension. This derivative can be thought of as a diagonal matrix, which is of course self adjoint.

### 2.4. Continuous versus discrete interpolation

For purposes of comparison, we include the continuous version of interpolation, “compose two functions” or equivalently “evaluate a function at a set of points”: *J* = ^f^(*I, φ*^-1^) ≐ *I*(*φ*^-1^).

#### Theorem 2.3.

*In the continuous case, the adjoint of the derivative of interpo-lation f (I, φ^-1^) = I(φ^-1^) with respect to its first and second argument is*

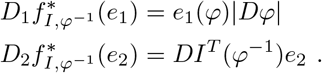

*When e_2_ is a vector, this denotes matrix multiplication.*

*Proof.* For the first argument, the operation is linear, so the derivative is just the operation itself.

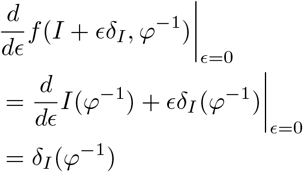

We evaluate the adjoint through a change of variables

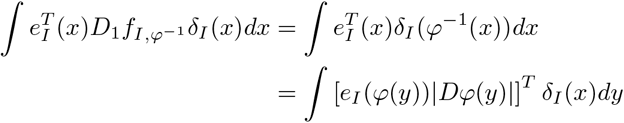

The operation is nonlinear in the second argument.

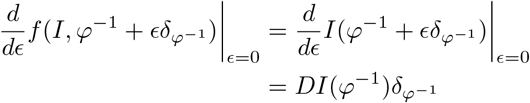

For clarity, we emphasize *DI*(*φ*^-1^) ≠ *D*[*I*(*φ*^-1^)].

The adjoint is calculated by

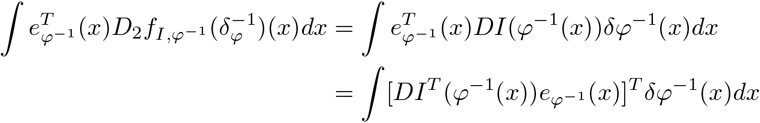

These results are “similar” to, but distinct from, those derived in the discrete section. For the first argument, when |*D_φ_*| is large, in the continuous setting we have a large multiplicatiave weight, whereas in the discrete setting we have many voxels in the accumulation. A small |*D_φ_*| implies a small weight in the continuous setting, but a sparse set of voxels in the discrete setting. The latter is an important fundamental difference between the two, when the transformation *φ* is contractive, many image voxels do not get sampled at all. The continuous approach artificially assigns them a nonzero gradient with respect to the cost.

For the second argument, the continuous formulation gives a derivative, while the discrete formulation gives a discrete forwards difference. Many existing registration implementations such as ITK [15] have used a centered difference, because its truncation error is accurate to second order. Other publications have explicitly [16] or implicitly [17] avoided considerations of discrete interpolation or derivatives.

### 2.5 The registration algorithm

Unknown parameters *v,w,S,R,θ* are estimated using the EM algorithm as described in [12]. In the E step, *W* is calculated as the posterior probability that each pixel corresponds to a deformation of the atlas (as opposed to an artifact or missing tissue). In the M step, all parameters are updated using gradient descent, calculating gradients via backpropagation as described above. For velocity fields *v, w*, we use the Hilbert gradient rather than the *L*_2_ gradient [17]. We force *S* (*R*) to be positive definite (rigid) by using the matrix log and exponential of a symmetric (antisymmetric) matrix. Gradient descent step sizes are fixed and their scale is chosen manually. More sophisticated optimization techniques, including replacing optimization with prediction[13], is the subject of future work.

## 3 Experiment and results

We demonstrate the performance of our algorithm by registering 2D slices to the 3D CCF. We apply this method to reconstructing a challenging dataset from the BICCN, which includes artifacts due to strong fluorescence in certain regions.

Figure 6 shows the result of our algorithm applied to BICCN fluorescence microscopy data from the laboratory of Hongwei Dong (USC), publicly available through https://biccn.org/data. The top row shows several slices of our observed data, while the second row shows our template transformed to match each slice using the generative model described above. The bottom row shows our algorithm’s identification of pixels that correspond to the atlas (red), versus missing data (blue) or artifact (green). Fig. 7 shows the cost function gradient, backpropagated into the 3D space of the atlas. One can see the effect of sparse sampling here. Many of the pixels do not contribute to the cost function, and their associated gradient is zero.

**Figure 6:**
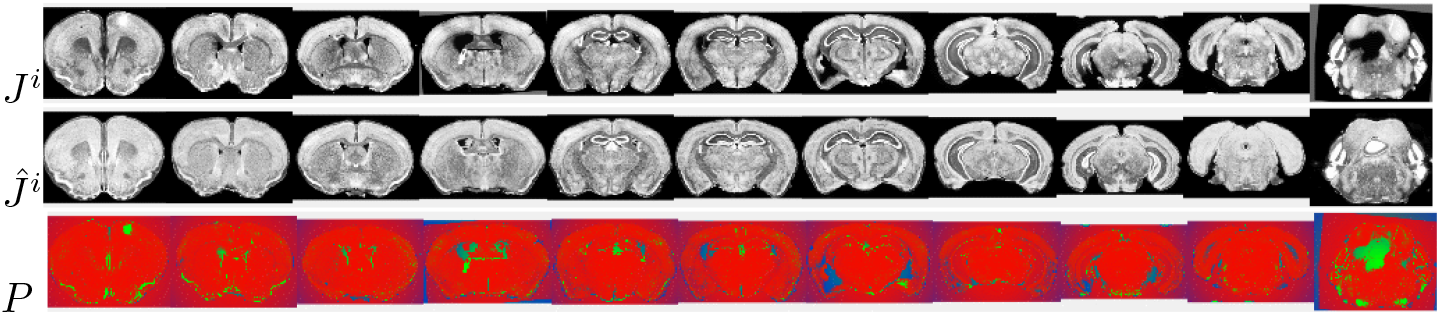
Top: a selection of 11 slices *J^i^* from the BICCN dataset. Second row: Corresponding deformed atlas slices with intensity transformation 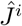. Bottom row: Estimates of posterior probability of missing tissue (blue), artifact (green), or data corresponding to the atlas (red). Note that alignment is poor toward the posterior cerebellum where the Allen atlas abruptly ends.

**Figure 7:**
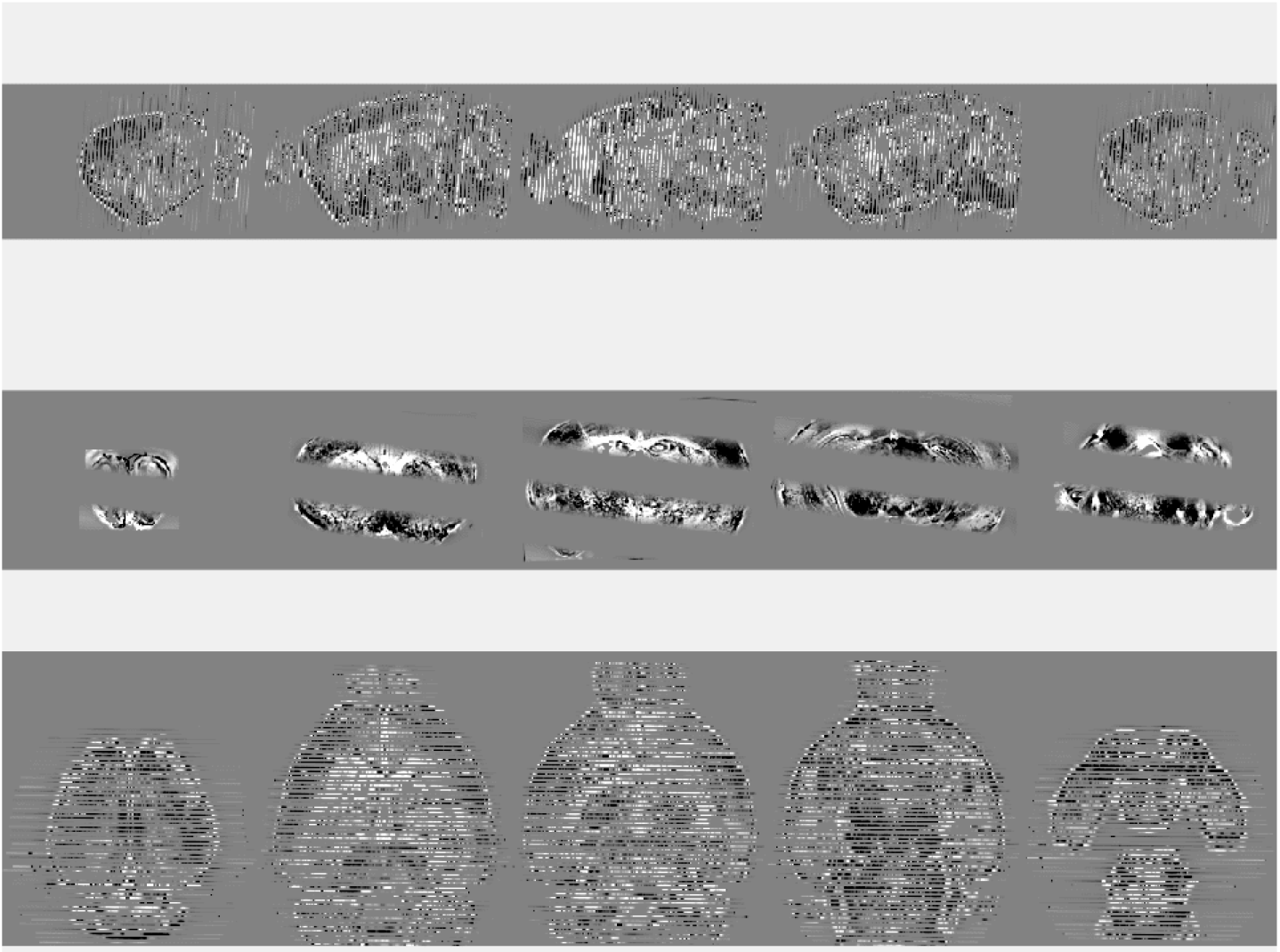
We show error backpropagated into 3D deformed atlas coordinates with five slices in sagittal (top), coronal (middle), and axial (bottom) planes. Grey implies little error, while black or white imply more error. The sparse sampling is evident with no error in between sectioning planes. Note that regions identified as artifact or missing data have a small weight on their error.

## 4 Discussion

In this work we formulated a 3D to 2D registration algorithm based of a generative statistical model of imaging, that is capable of functioning accurately and robustly in the presence of sparse slicing, artifacts, or missing tissue. Because this model involves joint optimization over a sequence of transformations, we used a computation graph to coordinate each operation and the adjoint of its gradient. We used this as an opportunity to analyze the effect of discretization in image registration, as opposed to a continuous formulation. We demonstrated the performance of this algorithm using simulated data and fluorescence microscopy data that is part of the BICCN.

A common challenge of deformable image registration is the presence of local minima in the objective function, and the resulting sensitivity to initialization of of the transformation parameters. The EM algorithm used here can make this problem more severe. We use several strategies to overcome this challenge, including a multiresolution approach (registering low resolution images with smoother deformations can have as an initialization to higher resolution), and a reasonable initial slice alignment by detecting tissue center of mass.

One limitation of this work is that we considered only mouse imaging data. The community commonly uses a variety of model organisms, and the BICCN intends to pursue nonhuman primate and human data. While this algorithm can easily be applied to other datasets, more work is needed to choose and validate appropriate parameters in each case.

In addition to analyzing model organisms for neuroscience research, we are applying these techniques to serial sections in human digital pathology. Three dimensional analysis of tissue specimens will be important for modern pathology departments. Unlike neuroscience experiments, acquiring human specimens cannot simply be replicated, and designing algorithms that function robustly in the presence of staining variability and missing or damaged tissue is essential.

## A Transformation parameterization

Affine transformations with linear part *A^i^* ∈ ℝ^3×3^ and translation part *b^i^* ∈ ℝ^3^ are decomposed using a polar factorization. this gives a positive symmetric definite scale matrix *S* used for all slices, and a rigid transformation *R^i^, b^i^* that is slice specific (see Fig. 2). These transformations are parameterized using the matrix group exponential with *S* = exp(*s*) where s is a symmeric matrix, and *R^i^* = exp(*r^i^*) where *r^i^* is an antisymmetric matrix.

Deformations are parameterized using the Riemannian exponential of a smooth initial velocity field *υ*_0_. Smoothness is ensured by modeling velocity fields as belonging to a Hilbert space *V* with inner product defined through a linear operator *L* [18]

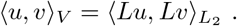

Diffeomorphisms are constructed via the flows

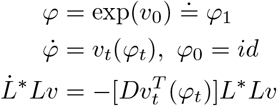

where * denotes the adjoint and *L*L_υ_* denotes the Eulerian momentum [19]. Computationally, we optimize over time varying velocity fields, allowing the Riemannian exponential trajectories to be approached at convergence.

